# Ant parasitoidism in checkered beetles: *Phyllobaenus obscurus* developing inside intact cocoons of two species of the *Ectatomma ruidum* species complex

**DOI:** 10.1101/2025.10.10.681666

**Authors:** Gabriela Pérez-Lachaud, Chantal Poteaux, John M. Leavengood, Jean-Paul Lachaud

## Abstract

Known parasitoids of ants include species from several families of flies, wasps, strepsipterans, nematodes, and mites. Curiously, while myrmecophily is heavily biased towards Coleoptera, one of the most diverse and speciose insect orders, no beetles specialized as parasitoids of ants have been recorded, although the parasitoid habit has evolved at least 13 times within this order. Here we report on observations that strongly suggest that a checkered beetle species behaves as a parasitoid of ant brood. A total of 146 colonies or part of colonies of three species of the *Ectatomma ruidum* species complex (*E. ruidum* sp. 2, 3, 4) were excavated in several sites along the Pacific coastal plains of Oaxaca, Mexico, during three collecting campaigns (2015-2017). Overall, 11060 adults, 5795 cocoons and 2185 larvae were examined. Upon dissection, four intact cocoons contained ant prepupae/pupae parasitized by characteristic campodeiform beetle larvae (prognathous head, three pairs of segmented legs on thorax, no prolegs, body with sparse but long pubescence), and a fifth cocoon presented a round exit hole. An active, pink-colored larva, emerged from a cocoon in 2015, was reared to the adult stage and could be identified as *Phyllobaenus obscurus* (Gorham) (Cleridae). Second and third instar larvae were found inside intact cocoons of two species: *E. ruidum* sp. 3 and sp. 4. The prevalence of parasitism is extremely low: less than 0.6% of cocoons available in each *Ectatomma* host populations. Predatory during both adult and larval stages, checkered beetles are broadly known as predators of wood-boring and cone-boring beetles, and some species are facultative parasitoids of solitary bees or wasps or, very rarely, specialized in predating social insects. We assert that the novel discovery of clerid-ant brood parasitoidism within the subterranean host colony deviates yet further from any known clerid adaptation to date.

## Introduction

Many invertebrates have adopted a predatory or parasitoid life strategy and play an important role in biological control [1–3]. A parasitoid is an organism in which the immature stages develop on or within a single organism (the host), with this feeding activity eventually resulting in the death or sterilization of the host [4,5], whereas adults are free-living. A diverse array of parasitoids specializes in attacking adult ants or their brood. Known ant parasitoids include species of several insect orders and families: flies, twisted-wing parasites, and parasitic wasps [6–11], two families of Nematoda [12], and one mite family [13–15]. Ant parasitoids are highly specialized symbionts that display a variety of morphological, chemical, and behavioral strategies for accessing and successfully parasitizing their hosts [13,16,17]. For example, species in the genera *Elasmosoma* Ruthe and *Neoneurus* Haliday (Hymenoptera, Braconidae) that parasitize adult ants have morphological adaptations for grasping workers and securing oviposition (e.g., tarsal modifications, raptorial legs) and rely on rapidity to oviposit directly into the ant body [18]. Wasps in the Eucharitidae family lay eggs away from their hosts, in or on plant substrate; they possess a very mobile first-instar larva, termed *planidium,* which is responsible for accessing the hosts through phoresis on foraging ants or attached to prey [19–23]. Chemical volatiles of glandular origin used by ants as pheromones are co-opted by ant-decapitating flies (Phoridae) to locate potential hosts [24]. Once in the vicinity, phorid females rely on short-range chemical signals, such as cuticular hydrocarbons, to locate and select potential hosts [25].

According to Eggleton and Belshaw [4,26], the parasitoid habit has evolved independently at least 13 times in Coleoptera. Beetles do not have piercing ovipositors and the poor flying ability of the adult contrasts with the mobility of the larval stages [4,26]. As in the case of eucharitid wasps and Strepsiptera, known species of Coleoptera with a parasitoid lifestyle lay eggs away from the hosts and possess phoretic first-instar larvae that are responsible for locating and selecting suitable hosts (Meloidae, Ripiphoridae, some Cleridae [4,19]). The phoretic larvae attach to the adults of their hosts either on living plants, dead wood or soil. Most parasitoid coleopteran families appear to have evolved from ancestors associated with dead wood, with some shifting to attacking insects in the soil. Host orders of coleopteran parasitoids include Diplopoda, Thysanura, Blattaria, Thysanoptera, other Coleoptera, Hymenoptera (other than Formicidae), and Lepidoptera [4]. Compared with parasitoid wasps and flies, their host taxonomic diversity is rather low, suggesting that other factors may constrain the host range of coleopteran parasitoids [4]. Contradictorily, evolution of myrmecophily is heavily biased towards the Coleoptera. The nature of colony exploitation by myrmecophilous beetles varies significantly, with species ranging from scavengers and refuse dwellers to tolerated or highly integrated guests [27]. Some adult myrmecophilous beetles, as in the genus *Cremastocheilus* Knoch [28] or in various Paussinae species [29], are predatory on ant brood and exhibit various strategies to locate and exploit their prey. Some species, such as *Myrmedonota xipe* Mathis & Eldredge, can even use specialized parasitoid host location cues to selectively prey on *Azteca sericeasur* Longino workers that have been previously parasitized by phorid flies [30]. However, to the best of our knowledge, no species of beetle has ever been recorded as developing as a parasitoid of adult ants or their brood. Here we report on observations that strongly suggest that a species of Cleridae develops as a parasitoid of ant brood.

Cleridae is a family of about 4710 described species (R. Gerstmeier, personal communication). Predatory during both adult and larval stages, these beetles are broadly known as predators of wood-boring and cone-boring beetles (e.g., bark beetles, longhorn beetles, powderpost beetles) and are often observed in the galleries of their prey [19,31]. In North America, this most accurately applies to the genera *Chariessa* Perty*, Cregya* LeConte*, Cymatodera* Gray*, Enoclerus* Gahan*, Madoniella* Pic*, Monophylla* Spinola*, Neorthopleura* Barr*, Pelonium* Spinola*, Priocera* Kirby*, Pyticeroides* Kuwert*, Tarsostenus* Spinola, and *Thanasimus* Latreille, recorded in forestry studies and occasional anecdotal observations during sampling [31]. However, several clerid genera are known to feed on gall-makers and their parasitoids (e.g., *Cymatodera*, *Neohydnocera* Leavengood, *Phyllobaenus* Dejean, *Placopterus* Wolcott), on pollen as a primary or secondary protein source (some Clerinae, Korynetinae, Epiclininae), on grasshopper egg pods (*Trichodes* Herbst, *Aulicus* Spinola), on non-gallmaking caterpillars (e.g., *Aulicus, Phyllobaenus*), in the nest cells of larval bees and wasps (e.g., *Cymatodera, Lecontella* Wolcott & Chapin, *Phyllobaenus, Placopterus*, *Trichodes*), on insects associated with carrion (*Necrobia* Latreille) and stored food products and their pests (*Necrobia*) [31,32]. Many species exhibit several different larval feeding strategies, which all may appear to be specialized behaviors (see Table 1). For example, the larvae of *Placopterus thoracicus* (Olivier), which is strongly associated with wood-boring beetles, have been found inside insect galls, and the adults have been reared from cells of crabronid and pemphredonid wasps [48]. Similarly, in *Neohydnocera longicollis* (Ziegler), formerly referred to as *Hydnocera longicollis* Ziegler and *Isohydnocera curtipennis* (Newman), adults have been reported emerging from the galls of three different hosts in three different insect orders (Diptera, Hymenoptera and Lepidoptera) [38,45,48,56,57]. On the other hand, various species, mainly from the genus *Trichodes*, develop as cleptoparasitoids (development at the expense of a single host organism by at least partial usurpation of its food supply, resulting in the death of the host [4]), and many other species have been reared from galls or nest cells of Hymenoptera while also exhibiting other predatory behaviors [31,32,68] (see Table 1). In general, however, the natural histories of most species in these genera are poorly documented.

**Table 1.**
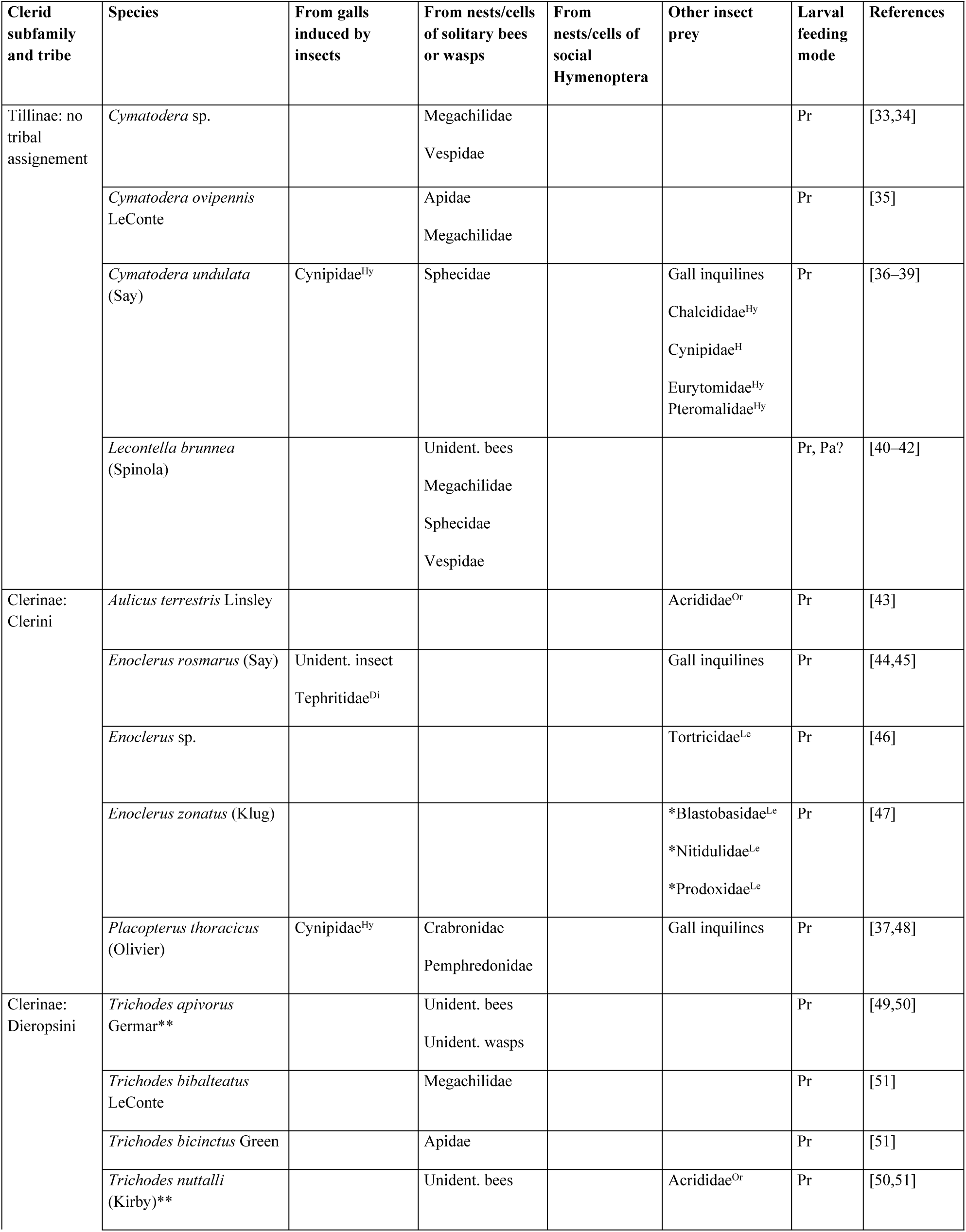
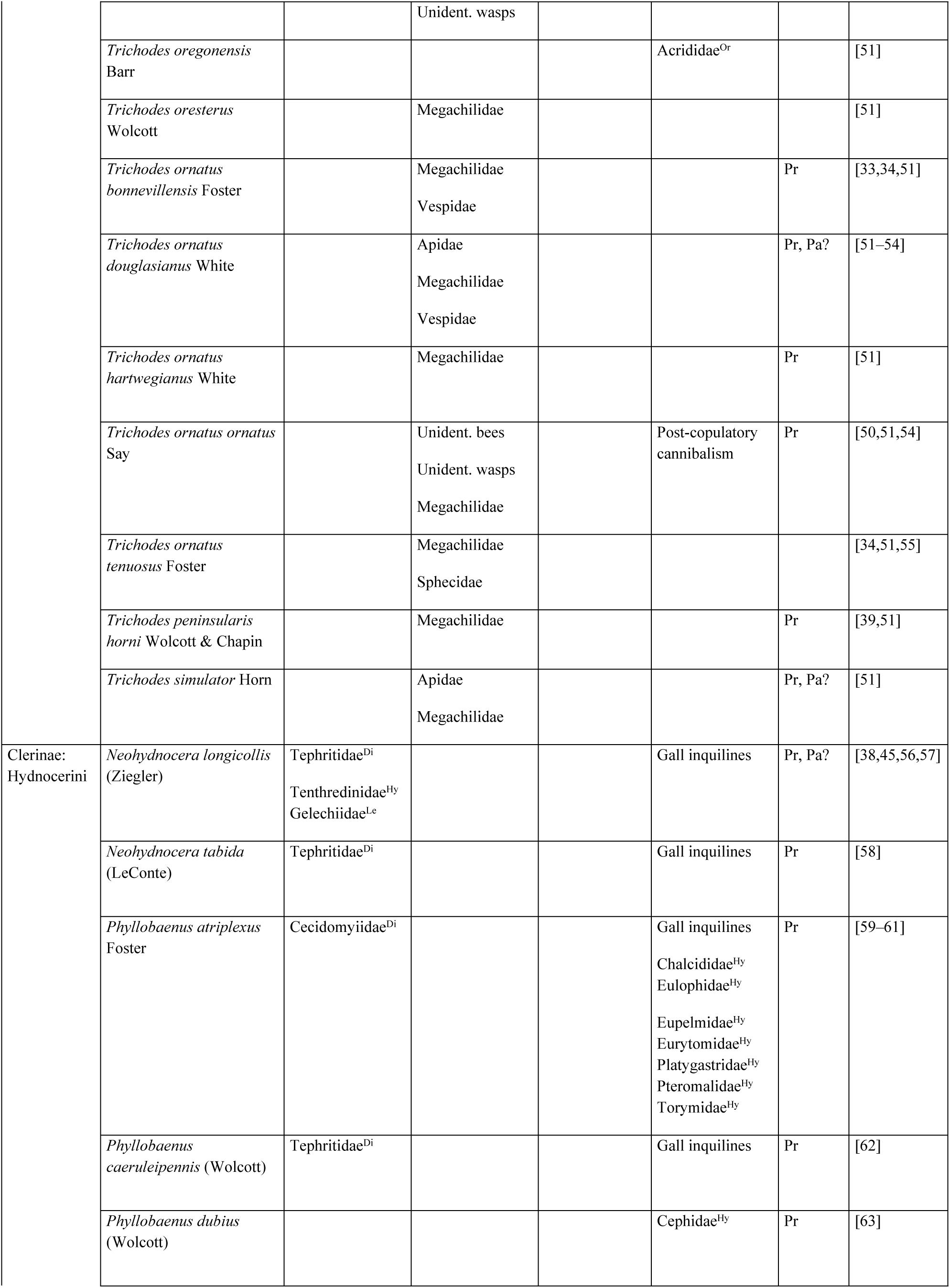
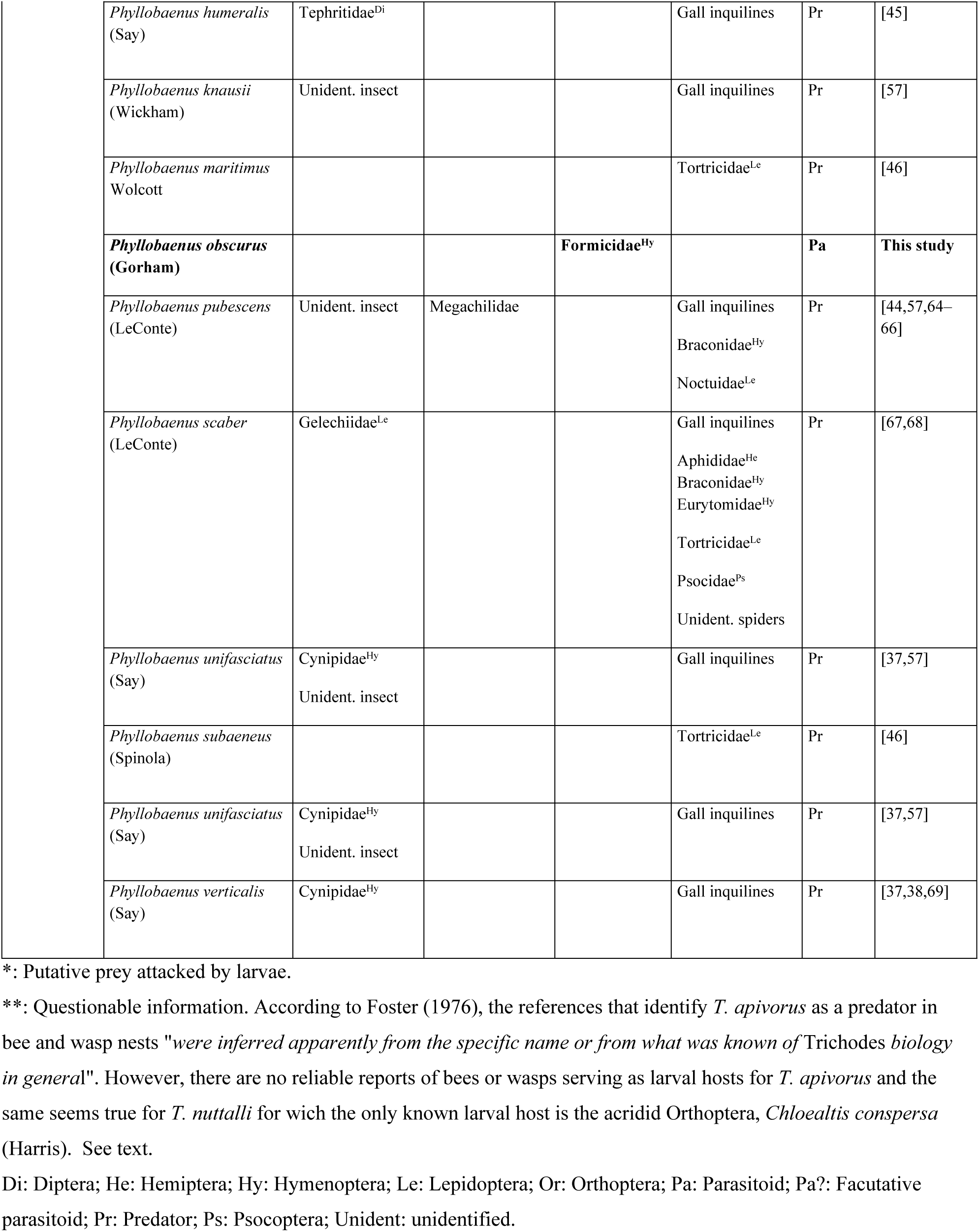
Summary of known rearing records among North American Cleridae and larval feeding mode on insects other than wood, stem and twig borers.

In Europe, the association between *Trichodes umbellatarum* Olivier and the larvae of various bee species borders parasitoidism and predation [70,71], some larvae developing upon a single host as facultative parasitoids while others need several hosts acting as predators. Similar shifts between predation and facultative parasitoidism have also been reported in the New World for some species such as *Lecontella brunnea* (Spinola), *Trichodes ornatus douglasianus* White, *T. simulator* Horn, and *N. longicollis* [42,51,52,57] (Table 1). However, no clerid species associated with hymenopterans can compare to the unique case of *Tarsobaenus letourneauae* Leavengood, Pinkerton & Rifkind and *T. piper* Leavengood, Pinkerton & Rifkind which exhibit a remarkable symbiotic relationship with ants and their host plants [72]. Larvae of both *Tarsobaenus* species have been reported inhabiting the petiole chambers of the Piperaceae *Piper cenocladum* Casimir de Candolle, and *P. obliquum* Ruiz & Pavón, respectively, which host the myrmicine ant *Pheidole bicornis* Forel, and feeding either on the ant brood when ants are present or on the plant-produced food bodies when ants are absent; remarkably, translocation experiments showed that, in the absence of ants, the clerid larva induced food-body production too [73]. As intriguing a case this parasitism of an ant-plant mutualism presents, we assert that the novel discovery of clerid-ant brood parasitism within *Ectatomma* colonies is without precedent among all previously known clerid adaptations to date. Here we report on the first recorded instance of a checkered beetle developing as a parasitoid of the ant brood which also represents the first primary parasitoid of ants from the order Coleoptera. We also summarize all known rearing records among North American Cleridae and conclude by discussing the evolutionary transitions and diversification of larval feeding habits within Cleridae.

## Material and methods

### The ant hosts

The neotropical ant *Ectatomma ruidum* (Roger) has historically been considered as a widely distributed ant taxon throughout the Neotropics, from Mexico to Brazil [74–77], readily visually recognized. These ants have been subject of intense investigation into colony organization, communication, etc., serving as a species model system for several studies including research on intraspecific cleptobiosis, a very rare behavioral trait [78–80]. Recent studies have revealed that this taxon represents indeed a species complex [77,81–84] including at least 6 putative species, which can be separated based on molecular, chemical and acoustic signatures, and on behavioral traits, but with a considerably conserved morphology which has prevented species description. Two species have a wide distribution, from Mexico to northern Argentina (*E. ruidum* sp. 1 and *E. ruidum* sp. 2), and four other species are apparently restricted to few localities in southern Mexico, along the coast of Oaxaca [83–84].

These ants occur in a wide range of habitats in the Neotropics, including plantations and conserved forests, from sea level up to 1600 m [85,86]. Colonies of these species are monodomous with one single exception [87]. Typically, *E. ruidum* nests have a single entrance about 3–4 mm wide dug in the ground [88,89]. In Chiapas, southern Mexico, colonies are made up of 50 to 200 individuals [90] and colony density can be very high locally (up to 11,200 nests per hectare [91]). These ants have a very generalist diet, and their foraging activity is mainly diurnal [89]. In general, colonies are monogynous, but in some regions they are facultatively polygynous with size-dimorphic queens (macro and microgynes) [92,93]. The variability in such traits has long suggested the existence of a complex of species.

### Study site and ant sampling

Beetle larvae were collected along with the ants as part of a larger project aimed at characterizing/unveiling cryptic species in the *E. ruidum* complex and assessing the diversity of some of their already recorded parasitoids (parasitic Hymenoptera, Diptera, and Nematodes, see [94]). Colonies or part of colonies were collected in several sites during three collecting campaigns (November 2015, October 2016 and October/November 2017) in the Pacific Coastal Plain of Oaxaca, Mexico (Fig 1, Table 2). Climate is typical of the Pacific watershed lowlands in southern Mexico, i.e., warm sub-humid with precipitation concentrated in the summer months (Aw0 [95]); mean annual temperature is 22-28 °C; mean annual precipitation is ca. 900 mm. The dominant plant communities in the lowlands (50–450 m asl) are classified as seasonally dry tropical forest [96].

**Fig 1.**
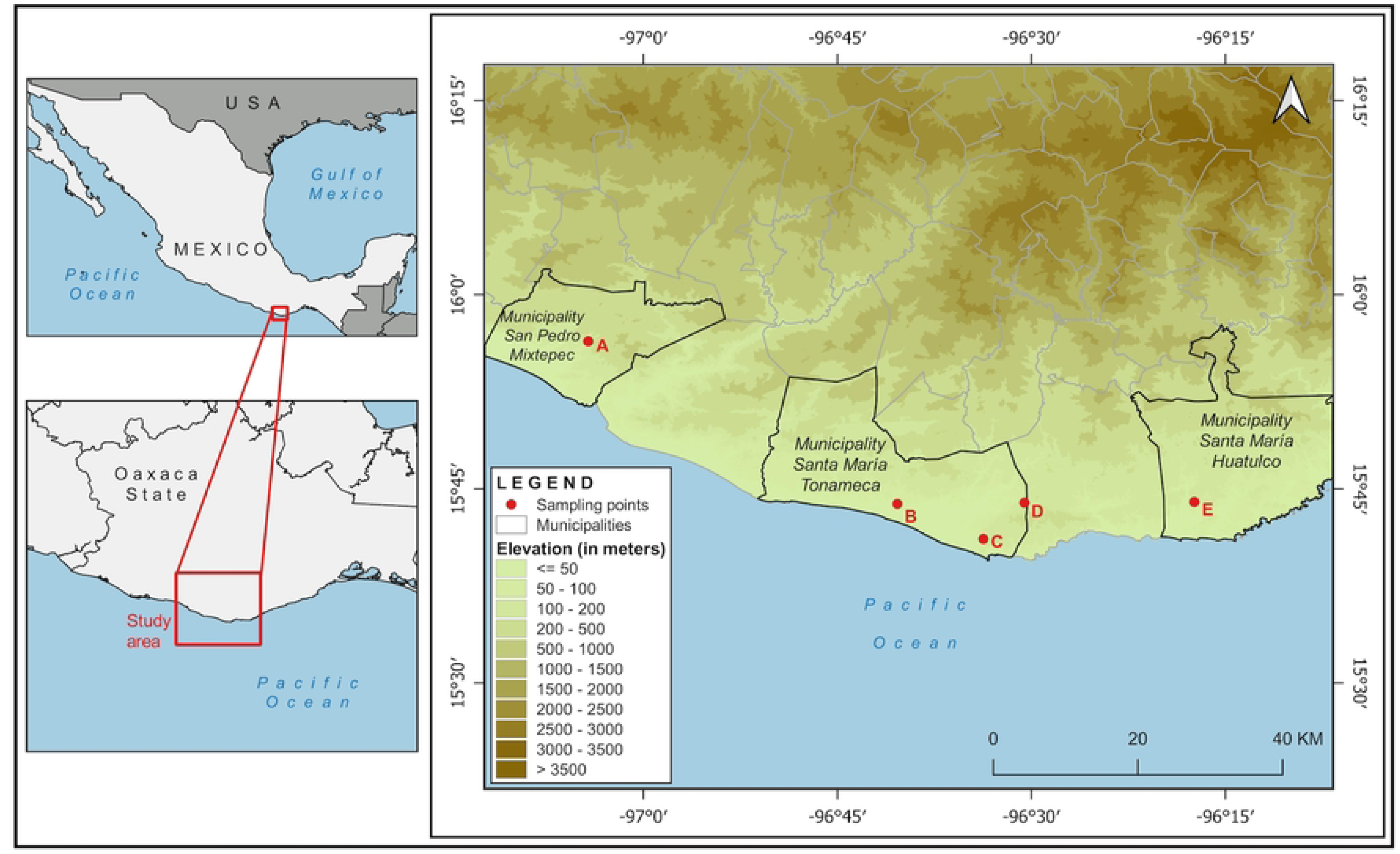
Location of the study sites in the coastal plain of Oaxaca, Mexico. A) Ranchería Yerba Santa; B) Piedras negras; C) El Zapotal (Mazunte); D) Puente Cuatode; E) Bajos de Coyula. Map credits: Holger Weissenberger.

**Table 2.** Summary of the collection sites in the Oaxaca coast (Mexico) and number of colonies of the different *Ectatomma ruidum* species sampled during 2015-2017.

| *Ectatomma ruidum*<br>species *sensu* Meza-Lázaro et al. [82] | Site | GPS coordinates | Altitud<br>(m asl) | Nb. of sampled colonies<br>(colonies with cocoons) |  |  |
| --- | --- | --- | --- | --- | --- | --- |
|  |  |  |  | 2015 | 2016 | 2017 |
| *E. ruidum* sp. 2 | Bajos de Coyula | 15°43'58.8" N, 96°17'24" W | 125 | 5 (4) | 28 (14) | - |
| *E. ruidum* sp. 3 | Yerba Santa | 15°56'28" N, 97°04'27" W | 156 | 4 (4) | 24 (18) | - |
| *E. ruidum* sp. 4 | Piedras Negras | 15°43'49.7" N, 96°40'21.7" W | 18 | - | 22 (13) | - |
| *E. ruidum* sp. 4 | El Zapotal<br>(Mazunte) | 15°41'7.4" N, 96°33'42.5" W | 42 | 4 (4) | 22(9) | 15 (15) |
| *E. ruidum* sp. 4 | Puente Cuatode | 15°43'55.2" N, 96°0'32.4" W | 89 | - | 22 (17) | - |
|  |  |  |  | 13 (12) | 118 (71) | 15 (15) |

At each site, the entrances of ant nests were detected using cookies as bait and following foraging ants back to their nest. Colonies were excavated and all the individuals present in the nest chambers were collected in plastic bags and transported to the field lab for initial revision. Cocoons of each colony were kept in Petri dishes for a couple of weeks to wait for parasitoid emergence. The rest of the material was preserved in 96° alcohol and revised and counted later under a stereomicroscope. The larvae were thoroughly examined for the presence of any *planidium* or evidence of parasitoid attack (presence of scars evidencing a previous unsuccessful attack or signs of endoparasite presence, see [9,11]). The cocoons were carefully dissected and their contents examined. The identity, number and developmental stage of the parasitoids were recorded. Adult ants were also closely examined for the presence of potential ecto- or endoparasites (e.g., phorid flies, strepsipterans, mites, nematodes, or *planidia*) attached to their body.

### Parasitoid identification

The initial assignment of wasps was performed using morphological characters and available keys [97]. Beetles were identified by one of us (JML). Voucher specimens of ants and parasitoids were deposited in the Formicidae and Arthropoda collections of El Colegio de la Frontera Sur-Chetumal (ECO-CH-F and ECO-CH-AR, respectively). Field sampling complied with the current laws of Mexico (collection permit FAUT-0277 from Secretaría del Medio Ambiente y Recursos Naturales - Dirección General de Vida Silvestre granted to GP-L).

## Results

A total of 146 colonies or part of colonies of three species of the *Ectatomma ruidum* complex (*E. ruidum* sp. 2, sp. 3, and sp. 4 *sensu* Meza-Lázaro et al. [82]) were sampled. Cocoons and/or larvae were present in a total of 98 excavated colonies or part of colonies (Tables 2, 3, Table S1). Overall, 11060 adults, 5795 cocoons and 2185 larvae were examined.

**Table 3.** Infestation rate of *Ectatomma ruidum* spp. cocoons by *Phyllobaenus obscurus*.

| Year of collection | Species | Locality | Nb. of colonies with cocoons | Total nb. of cocoons | Total nb. of larvae | % of cocoon infestation (exact proportion) |
| --- | --- | --- | --- | --- | --- | --- |
| 2015 | *E. ruidum* sp. 2 | Bajos de Coyula | 4 | 111 | 2 | 0 |
|  | *E. ruidum* sp. 3 | Yerba Santa | 4 | 172 | 72 | 0.6 (1/172) |
|  | *E. ruidum* sp. 4 | El Zapotal | 4 | 233 | 87 | 0 |
| 2016 | *E. ruidum* sp. 2 | Bajos de Coyula | 14 | 487 | 74 | 0 |
|  | *E. ruidum* sp. 3 | Yerba Santa | 18 | 1285 | 794 | 0 |
|  | *E. ruidum* sp. 4 | Puente Cuatode | 17 | 1229 | 371 | 0.3 (4/1229) |
|  |  | El Zapotal | 9 | 717 | 116 | 0 |
|  |  | Piedras Negras | 13 | 490 | 191 | 0 |
| 2017 | *E. ruidum* sp. 4 | El Zapotal | 15 | 1071 | 478 | 0 |
| Total |  |  | 98 | 5795 | 2185 | < 0.1 (5/5795) |

Workers were parasitized by nematodes (36 cases). The larvae were not parasitized, whilst at dissection eight cocoons were found parasitized by eucharitid wasps belonging to an undetermined species from the *Kapala* Cameron clade, and five cocoons were attacked by beetle larvae (Table 3). In 2015, 12 of the 13 colonies excavated had cocoons. From a total of 516 cocoons maintained in observation, a single, pink-colored larva emerged 10 days later (December 2) from cocoons of a colony of *E. ruidum* sp. 3 collected at Ranchería Yerba Santa (representing 0.6 % of cocoons of this species, Fig 2A, Video S1). The cocoon from which the larva had emerged showed signs of a single round exit hole. When it was dissected, the ant host pupa was found beheaded and only its empty exoskeleton remained (Fig 2B). The larva was isolated in a glass vial along with some fabric to help pupation. An adult belonging to a species of Cleridae (Insecta: Coleoptera) was reared in early January and could be identified as a specimen of *Phyllobaenus obscurus* (Gorham) (Cleridae: Clerinae: Hydnocerini) (Fig 2C). During the 2016 campaign, five sites and 118 colonies were sampled (71 with cocoons). Four cocoons from *E. ruidum* sp. 4 collected at Puente Cuatode, were parasitized by beetle larvae or presented signs of attack (Table 3). One first- and two second-instar larvae were obtained (Figs 3A–3C), each one found inside an intact cocoon on an ant pupa. Another cocoon presented a round hole, suggesting a larva had already left the host to pupate elsewhere (Fig 3D) and the remains of the host pupa were found inside. During 2017 only one site was sampled (El Zapotal, Mazunte; 15 colonies of *E. ruidum* sp. 4) due to instable sociopolitical conditions and insecurity in the region. No beetle or parasitoid wasp was found during this collecting campaign. Whatever the campaign, eggs of the Cleridae were never obtained.

**Fig 2.**
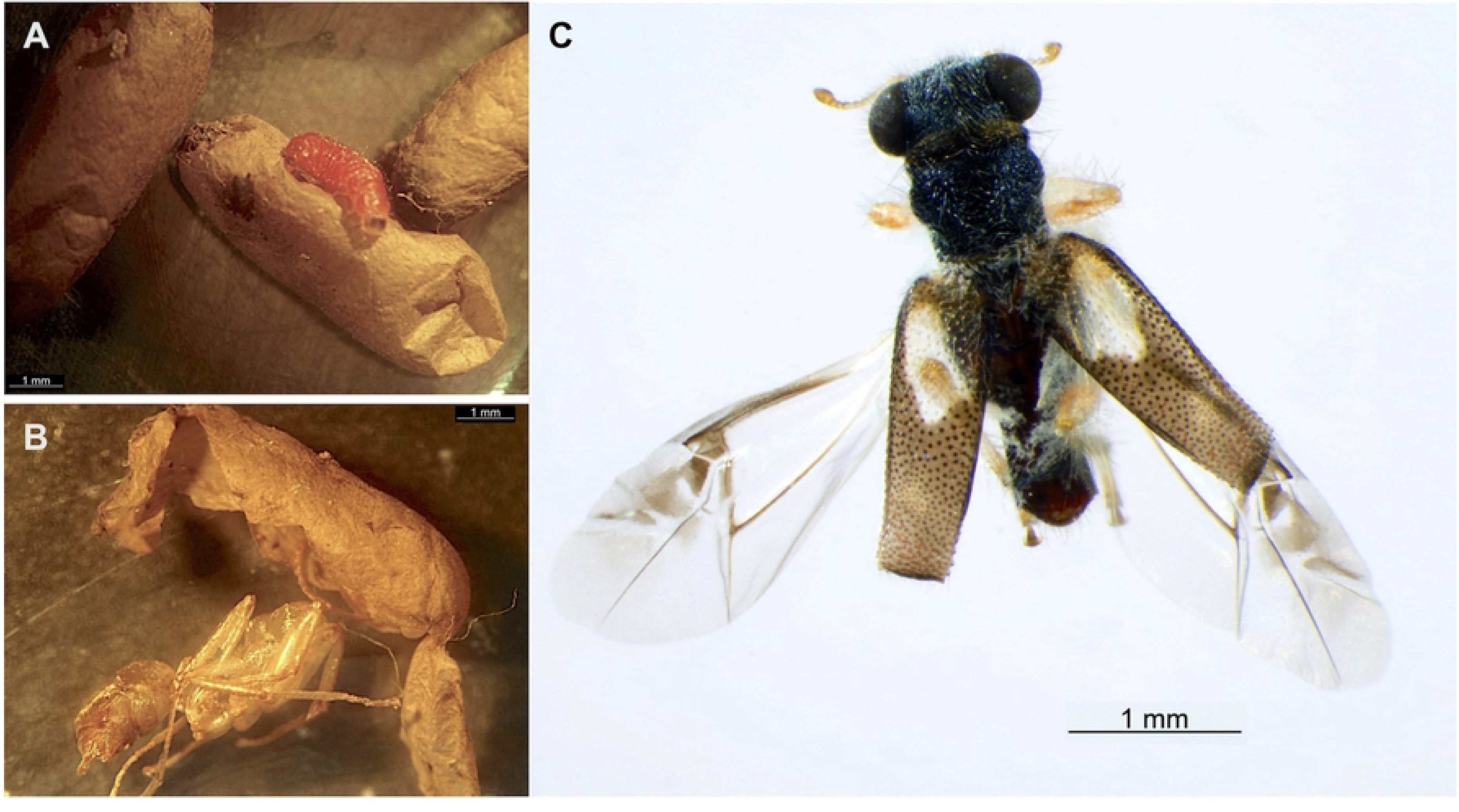
*Phyllobaenus obscurus* (Coleoptera: Cleridae). A) A third instar larvae that have just exited its host cocoon. B) Remains of an *Ectatomma* sp. 3 pupa from which a mature beetle larva has emerged (the dissected cocoon can be observed). C) *P. obscurus* adult reared from the larva that parasitized the pupae in (B). Photo credits: (A-B) G. Pérez-Lachaud, (C) Humberto Bahena-Basave.

**Fig 3.**
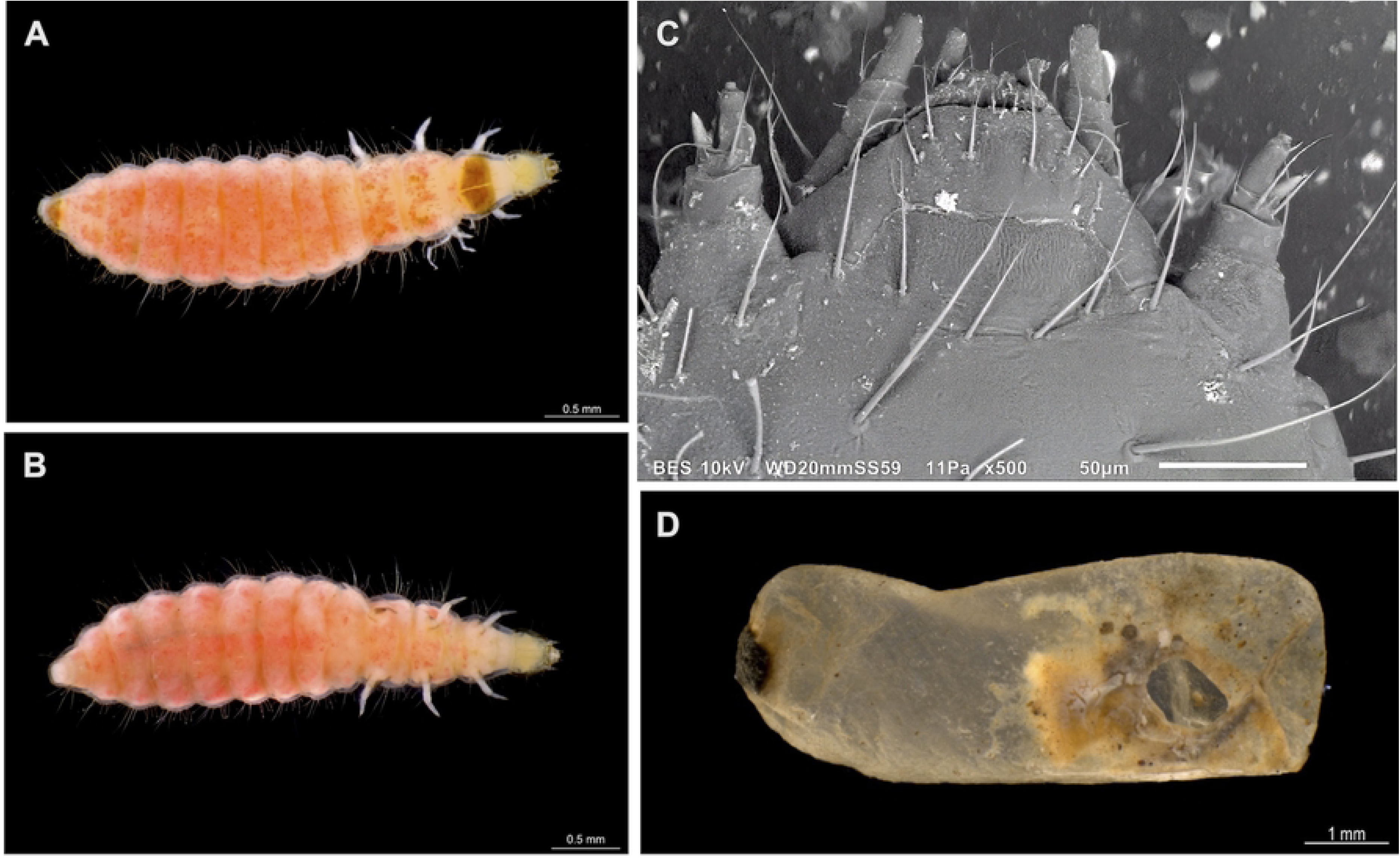
*Phyllobaenus obscurus* larva and remains of the host within the host cocoon. A) Third instar *P. obscurus* larva, dorsal view. B) Same, ventral view. C) Dorsal view of the head of a third instar larva, low vacuum SEM photography. D) An *Ectatomma* sp. 6 cocoon showing an exit hole after departing of a *P. obscurus* larva; the ant remains are visible. Photo credits: (A-B, D) Humberto Bahena-Basave; (C) G. Pérez-Lachaud and M. Elías-Gutiérrez.

## Discussion

The genus *Phyllobaenus*, the second largest Hydnocerini genus after *Callimerus* Gorham, comprises 114 described species [98–101]. Hydnocerini are considered general ant mimics, due either to their wasp-waisting color patterns, their elongate body shape (also sometimes creating a physical “waisting” effect), or their very movement patterns [102–104]. For example, *Neohydnocera aegra* (Newman) strikingly resembles a species of *Pseudomyrmex* Lund and has been found abundantly among these ants [105], and *Phyllobaenus cryptocerinus* (Gorham) is a reported mimic of *Cephalotes* Latreille ants (= *Cryptocerus* Latreille) [102].

Like many of its congeners, *P. obscurus* is an active clerid with a highly variable color pattern and body size (2-5 mm). This intraspecific plasticity can make species determination difficult as there are many similar Hydnocerini species [100]. This species is known from Mexico, Guatemala, Costa Rica, and Panama [106–109]. Araujo et al. [110] included Brazil based on an unvouchered record with no data nor depository presented from a personal database (see “Unverified Distributions” in [111]). Prior to this study it was known only from Durango, Morelos and Tamaulipas in Mexico [112,113]. The data presented herein for Oaxaca and an unpublished record from Chiapas ([Verbatim label data] 3000’, “El Chorreadero” by Hwy. 190,

1.25km E Tuxtla Gutierrez, under debris, gravel bank of stream in deep ravine, 25-XII-1972, H. Frania, FSCA, Florida State Collection of Arthropods) are new state records for Mexico. Despite the broad distribution of this rather common species, nothing was known about its natural history prior to our findings, and the larva and larval feeding habits were unknown.

Most specimens of *P. obscurus* have been collected by beating the foliage of live trees and occasionally shrubs (JML personal observations; accounts from numerous beetle collectors).

Specimen label data indicate few specimens collected with other methods (e.g., lights, fogging, Malaise trap) (Leavengood, unpublished data). Notably, label data for the Chiapas new state record for Mexico (see above) indicate an adult was caught on the ground and under debris, which is most unusual. In the Western Hemisphere, hydnocerines are not observed on the ground, but usually high in foliage (e.g., [104]) and there are no specimen records of *Phyllobaenus* collected in pitfall traps. Author JML has been presented two separate accounts of a *Phyllobaenus* specimen emerging from an ant hill in the ground. But no specimens were produced nor identifications made. So, with only one vouchered record (Chiapas, FSCA), two unvouchered events (ant hill emergence observations), and the five brood parasitoid observations noted herein (recorded in different years, at two different places, and concerning two different ant species), any record of *Phyllobaenus* at ground level (or below) is highly unusual, yet all but one such record are specifically associated with ants. Furthermore, it should be noted that our records concern clerid larvae which were sampled inside ant nests, in deep chambers.

Myrmecophilous taxa are not scattered randomly across the arthropod tree of life. Rather, their evolution is strongly biased to certain groups and, among insects, towards the Coleoptera which “*contains a large pool of lineages with the potential for transitioning to myrmecophily*” (Parker 2016). At least 33 of the beetle families, representing approximately one-fifth, have thus far been documented to include one or more taxa that exhibits close association with ant colonies [27]. Many of the most familiar beetle families contain at least one lineage that inhabits ant colonies, including the mainly aquatic beetle family Hydrophilidae [114]. Indeed, at least two species of *Sphaerocetum* Fikáček have been collected in mixed nests of *Camponotus* Mayr and *Crematogaster* Lund, and a larva of *S. arboreum* Fikáček et al. was collected in an ant nest along with the adults, implying the entire life cycle may take place inside the ant nest, suggesting that these beetles likely prey on ant larvae [114].

Ants and their nests are reliable, fixed resources in space, and potential targets that are available all year round [115]. In addition, as suggested by Parker [27], several ecological constraints and phenotypic attributes may predispose some taxa to undergo the evolutionary transition from free-living to myrmecophily which, in combination, could explain the myrmecophile beetle bias: i) the huge species richness of Coleoptera, ii) their ecological predisposition to encountering and exploring ant colonies, and iii) their possession of a major protective preadaptation in the form of elytra. Additionally, the size of the myrmecophile relative to the ant host species (small myrmecophile species/large host), a limuloid body form, and other protective morpho-anatomical adaptations (i.e., pedofossae) are traits that facilitate the integration of some intruders into colony life (see [13,116–118]).

Whether *P. obscurus* is a facultative or obligate parasitoid of the ant brood remains to be elucidated. Our finding of an unusual feeding mode for a checkered beetle is surprising but not entirely unexpected as other checkered beetles belonging to different tribes border the line between predator and parasitoid [42,51,52,57,70,71], likely depending on the relative size of their prey/host. It suggests that the parasitoid lifestyle may have evolved several times within the Cleridae. Notably, our record is also the first evidence of beetles parasitizing successfully the immature stages of ants within their subterranean nests, suggesting a major evolutionary shift in the biology of Hydnocerini. Our results further support a previously stated hypothesis about the evolution of ant parasitoidism in clades with predatory habits. As suggested for other unusual ant parasitoids such as the microdontine syrphid fly *Hypselosyrphus trigonus* Hull [119], the only parasitoid species among this clade of predators, it is likely that parasitoidism in *P. obscurus* evolved from a predatory ancestor. According to Godfray [120], the transition from a predatory to a parasitoid lifestyle may be straightforward since a predator that requires only a single prey item for its development is, by definition, a parasitoid.

There are still some gaps in knowledge regarding the biology of this clerid. Like other ant parasitoids such as eucharitid wasps of the genera *Kapala*, *Isomerala* Shipp, *Obeza* Heraty, or *Pseudochalcura* Ashmead [8], *P. obscurus* develops at the expense of host larvae/pupae under the protection of a cocoon, avoiding deadly interactions with their host ants. But, as our observations show, when larval development is completed, the larva exits the host cocoon to pupate. Clerids are known to commonly pupate near their prey (e.g., in galleries of their prey, or tree crevices nearby or inside the host pupal cell [57,58]), but the site of pupation of *P. obscurus* remains to be uncovered. Its high mobility and small size (only 2-5 mm for the adult) relative to the *Ectatomma* hosts (8-9 mm) may help avoiding ant aggressiveness. Infiltration into the ant nest is probably by phoresis of the mobile first-instar larva (*triungulin*) as occurs in other ant parasitoids which lay eggs away from the host [7,20,22]. Escape behaviors are an important part of myrmecophile strategies to adapt to their hosts, and the mobility of the adult may be responsible for leaving the natal nest unnoticed and the protective function of the elytra for coping with the host aggressiveness.

Although some checkered beetle species are considered of importance as potential biological control agents, there are very few rearing records for Cleridae in general, and data on their natural life history are scarce (Table 1). However, results of laboratory and field studies underscore the plasticity and adaptability in the food habits of Cleridae (e.g., [54] and references in Table 1). This deviating lineage (from predation) of foraging behavior towards parasitoidism reported here is likely to be a confirmation of the diversified feeding traits within the Cleridae.

## Acknowledgements

We are grateful to Ruby Meza-Lázaro, Alejandro Zaldívar-Riverón (UNAM, Mexico) and Carlos Santamaría (Universidad del Valle, Cali, Colombia) for their help during the 2016 field work; Humberto Bahena-Basave (Ecosur) for help with insect pictures, Holger Weissenberger (Ecosur) for help with the map, Manuel Elías-Gutiérrez (Ecosur) for help with SEM photography, and Roland Gerstmeier (Zoologische Staatssammlung München, Germany) for an up to date estimate of world clerid species.

## Supplementary material

**S1 Video**. A *Phyllobaenus obscurus* larva that has just emerged from its *Ectatomma ruidum* sp. 3 host cocoon.

**S1Table. Detailed collection data for the three species of the *Ectatomma ruidum* complex studied:** species concerned, location, date of collection, colony composition. µQ: number of microgynes.

